# A backward encoding approach to recover subcortical auditory activity

**DOI:** 10.1101/606251

**Authors:** Fabian Schmidt, Gianpaolo Demarchi, Florian Geyer, Nathan Weisz

**Author notes:** Address for correspondence: Fabian Schmidt, University of Salzburg, Centre for Cognitive Neuroscience, Hellbrunnerstr. 34, A-5020 SALZBURG, Austria/Europe.

## Abstract

Several subcortical nuclei along the auditory pathway are involved in the processing of sounds. One of the most commonly used methods of measuring the activity of these nuclei is the auditory brainstem response (ABR). Due to its low signal-to-noise ratio, ABR’s have to be derived by averaging over thousands of artificial sounds such as clicks or tone bursts. This approach cannot be easily applied to natural listening situations (e.g. speech, music), which limits auditory cognitive neuroscientific studies to investigate mostly cortical processes.

We propose that by training a backward encoding model to reconstruct evoked ABRs from high-density electrophysiological data, spatial filters can be tuned to auditory brainstem activity. Since these filters can be applied (i.e. generalized) to any other data set using the same spatial coverage, this could allow for the estimation of auditory brainstem activity from any continuous sensor level data. In this study, we established a proof-of-concept by using a backward encoding model generated using a click stimulation rate of 30 Hz to predict ABR activity recorded using EEG from an independent measurement using a stimulation rate of 9 Hz. We show that individually predicted and measured ABR’s are highly correlated (*r* ∼ 0.7). Importantly these predictions are stable even when applying the trained backward encoding model to a low number of trials, mimicking a situation with an unfavorable signal-to-noise ratio. Overall, this work lays the necessary foundation to use this approach in more interesting listening situations.

## 2. Introduction

Following mechanoelectrical transduction in the cochlea, neural activity first passes through several subcortical nuclei. Along the ascending auditory pathway these nuclei perform several fundamental computations relevant for sound processing (Romand & Ehret, 1997). However, an abundance of descending pathways exist in the auditory system involving brainstem and even cochlear structures, with a still weakly understood functional role (Terreros & Delano, 2015). With most of the findings stemming from animal research (for reviews, see Huffman & Henson, 1990; Suga & Ma, 2003; Terreros & Delano, 2015), the availability of tools to study neural activity along subcortical auditory regions in healthy human participants would be of outstanding value in broadening our understanding of their functional roles in normal and disordered hearing.

In humans, inference about subcortical auditory activity is usually drawn from the noninvasively measured auditory brainstem response (ABR) – a sequence of EEG evoked potentials that occur within the first 10 ms after acoustic stimulation. The recording consists of five to seven vertex positive waves (Wave I to VII; Jewett & Williston, 1971). By convention, each of these waves have been associated with activity in one or a few subcortical structures along the auditory pathway (Biacabe, Chevallier, Avan & Bonfils, 2001). As electrical signals also have magnetic properties an ABR can in principle also be captured using magnetoencephalography (MEG; Lütkenhöner, Lammertmann, Ross, & Pantev, 2000; Parkkonen, Fujiki, & Mäkelä, 2009). However, in practice this has been controversial, given the deep location of brainstem structures and the pronounced sensitivity of MEG to superficial sources (Hari & Salmelin, 2012). Yet, a recent study using concurrent intracerebral and MEG recordings shows that activity in deep subcortical areas (Amygdala, Hippocampus) can be disentangled from the surface MEG activity (Pizzo et al., 2019). In a pioneering study, Parkkonen et al. (2009) measured a magnetic ABR (mABR). Equivalent current dipole models offered plausible source localizations of the ABR components. While these results are promising, the general paradigm used to measure (m)ABR’s poses strong limitations: In particular due to the low signal-to-noise ratio of brainstem activity on M/EEG sensors, obtaining them usually requires averaging over thousands of simple sounds such as clicks or tone bursts to elicit a clear response. This is especially problematic for researchers interested in the role of brainstem regions in more naturalistic listening tasks (e.g. listening to continuous speech or music). This makes the availability of tools to further study these neural activity patterns increasingly important.

An important step in this direction was undertaken by Maddox and Lee (2018). By performing a regression between a rectified speech audio signal and a concurrent EEG recording they measured an ABR to continuous speech. They showed a high correlation between this speech-derived and a previously measured click-evoked ABR, suggesting that ABR’s can be derived from natural speech. However, these modeled ABR’s will only be possible under ongoing sound stimulations, with segments that furthermore are long enough to enable the computation of a temporal response function (TRF). Using a similar approach, by cross-correlating the fundamental frequency time-course of continuous speech with the recorded EEG signal, Forte, Etard and Reichenbach (2017) found a distinct peak at ∼9 ms. This evoked activity was modulated by attention, leading the authors to suggest a mechanism of selective attention at the level of the brainstem. Apart from likely not exclusively reflecting brainstem effects (see discussion), the approach still has the aforementioned limitation of only being applicable to paradigms using continuous stimulation.

Here, we propose a new approach to overcome this issue, that allows for modeling of auditory brainstem activity within scenarios where the signal-to-noise ratio is low. We used the signals obtained during a concurrent electric and magnetic ABR recording to generate channel weights that are tuned to putative auditory brainstem activity. These weights were computed by using the signal captured by the MEG sensors as regressors for a concurrent electric ABR recording. These weights could then be applied to MEG data recorded in an independent measurement to estimate auditory brainstem activity.

In this initial study, we attempt to establish the proof-of-concept that by training a model on a 30 Hz stimulation rate we can predict the dynamics of an ABR from an independent measurement using a different stimulation rate (9 Hz). By showing that the individually reconstructed and measured ABR’s are highly correlated even when parametrically reducing the signal-to-noise ratio, we lay the necessary foundation to generalize this approach to more interesting and complex listening situations.

## 3. Materials & Methods

### 3.1 Subjects

The data, collected from 14 healthy volunteers (10 males, *M* = 26.71 years; *SD* = 4.38 years) without reported psychiatric, neurological and hearing disorders, was analyzed for this study. The experimental protocol was approved by the ethics committee of the University of Salzburg and all participants gave written informed consent before the beginning of the experiment.

### 3.2 Stimuli and procedure

Participants listened to a total of 10000 clicks per condition at a sound pressure level of 60 dB. Stimuli were presented binaurally in two separate blocks at a rate of 30 Hz and a rate of 9 Hz. Each click stimulus had a duration of 80 μs. Auditory stimuli were presented inside the MEG helmet through a pneumatic system (SOUNDPixx, Vpixx technologies, Canada) with a stimulation delay of 16.5 ms estimated using a microphone (ER10C, Etymotic Research, US) and an oscilloscope (Rigol DS 1074Z). This delay was compensated by shifting the time axis during the data analysis accordingly. The measurement took in total about 6 minutes for the 30 Hz condition and about 20 minutes for the 9Hz condition. The experimental procedure was programmed using Psychophysics Toolbox 3 (Brainard, 1997) and custom built Matlab routines (Mathworks, Natick, USA).

### 3.3 Data acquisition

Magnetic brain activity was recorded using a 306-channel whole head MEG system (TRIUX, Elekta Oy, Finland) with a sampling rate of 10 kHz. The system consists of 204 planar gradiometers and 102 magnetometers. Before entering the magnetically shielded room (AK3B, Vakuumschmelze, Hanau, Germany), the head shape of each participant was acquired with >300 digitized points on the scalp, including fiducials (nasion, left and right pre-auricular points) with a Polhemus FASTRAK system (Polhemus, Vermont, USA). The auditory brainstem response (ABR) was measured with a single electrode located on FpZ based on the electrode placement of the international 10–20-System (Klem, Lüders, Jasper, & Elger, 1999). A ground electrode was placed on the forehead at midline and a reference on the clavicle bone of the participants.

### 3.4 Data analysis

#### 3.4.1 Preprocessing

The acquired data was Maxwell-filtered using a Signal Space Separation (SSS) algorithm (Taulu & Simola, 2006) implemented in the Maxfilter program (version 2.2.15) provided by the MEG manufacturer to remove external magnetic interference from the MEG signal and realign data to a common standard head position (-trans default Maxfilter parameter). As planar gradiometers are less sensitive to sources below the cortical surface than magnetometers (Vrba & Robinson, 2002) only magnetometer data was included in the following analysis. The Maxwell-filtered and continuous data was then further analyzed using the FieldTrip toolbox (Oostenveld, Fries, Maris, & Schoffelen, 2011) and custom built Matlab routines. First, the data was high-pass filtered at 1 Hz and low-pass filtered at 1000 Hz using a finite impulse response (FIR) filter (Kaiser window). For extracting physiological artefacts from the data, 50 independent components were calculated from the filtered data. Via visual inspection, the components showing eye-movements & heartbeats were removed from the data. On average 3 components were removed per subject (*SD* = 1). The data was then again high-pass filtered at 150 Hz and chunked into epochs of 50ms pre-/ & post-stimulus onset.

#### 3.4.2 Backward modeling

In order to reconstruct auditory brainstem activity from the MEG a backward encoding model was computed by using the signal captured by the MEG sensors (during the first 10ms) as regressors for a concurrent electric ABR recording. This was achieved by performing a ridge regression between the electric ABR and the data obtained from the magnetometers utilizing the mTRF toolbox (Crosse, Di Liberto, Bednar, & Lalor, 2016). A 5-fold cross-validation procedure was implemented and the regularization parameter (λ) that optimized the mapping between the EEG and MEG signal (for the 30 Hz stimulation rate condition), was selected based on a procedure suggested by Willmore & Smyth (2003). Instead of the usual trial-to-trial based cross validation approach, sub-averages (per 100 trials) were built to speed up processing and account for the low signal-to-noise ratio of brainstem activity. Afterwards the backward model was trained by utilizing the mTRF toolbox (Crosse et al., 2016) to perform a ridge regression between the signal captured by the MEG and EEG to generate channel weights for each training time point **(Figure 1, Backward modeling)**. By applying these weights to a new MEG measurement, a matrix with the time generalized “activation” for each testing/training time point is created **(Figure 1, Prediction Matrix)**. Analogous to classification based decoding approaches (King & Dehaene, 2014), we assume that this temporal generalization can be interpreted as the presence of an activation pattern during a training period (e.g. the pattern contributing to a specific ABR wave) at a corresponding testing time. While this approach could be flexibly applied to any other paradigm, we seeked to establish an important proof-of-principle for this approach: i.e. here we tested whether backward models obtained from magnetometers during one ABR measurement using a 30 Hz stimulation rate could correctly predict the ABR obtained from an independent measurement using a 9 Hz stimulation rate **(Figure 1, Evaluation)**. In order to compare the reconstructed with the measured signal, an overall correlation coefficient between both signals was computed for each subject individually.

**Figure 1:**
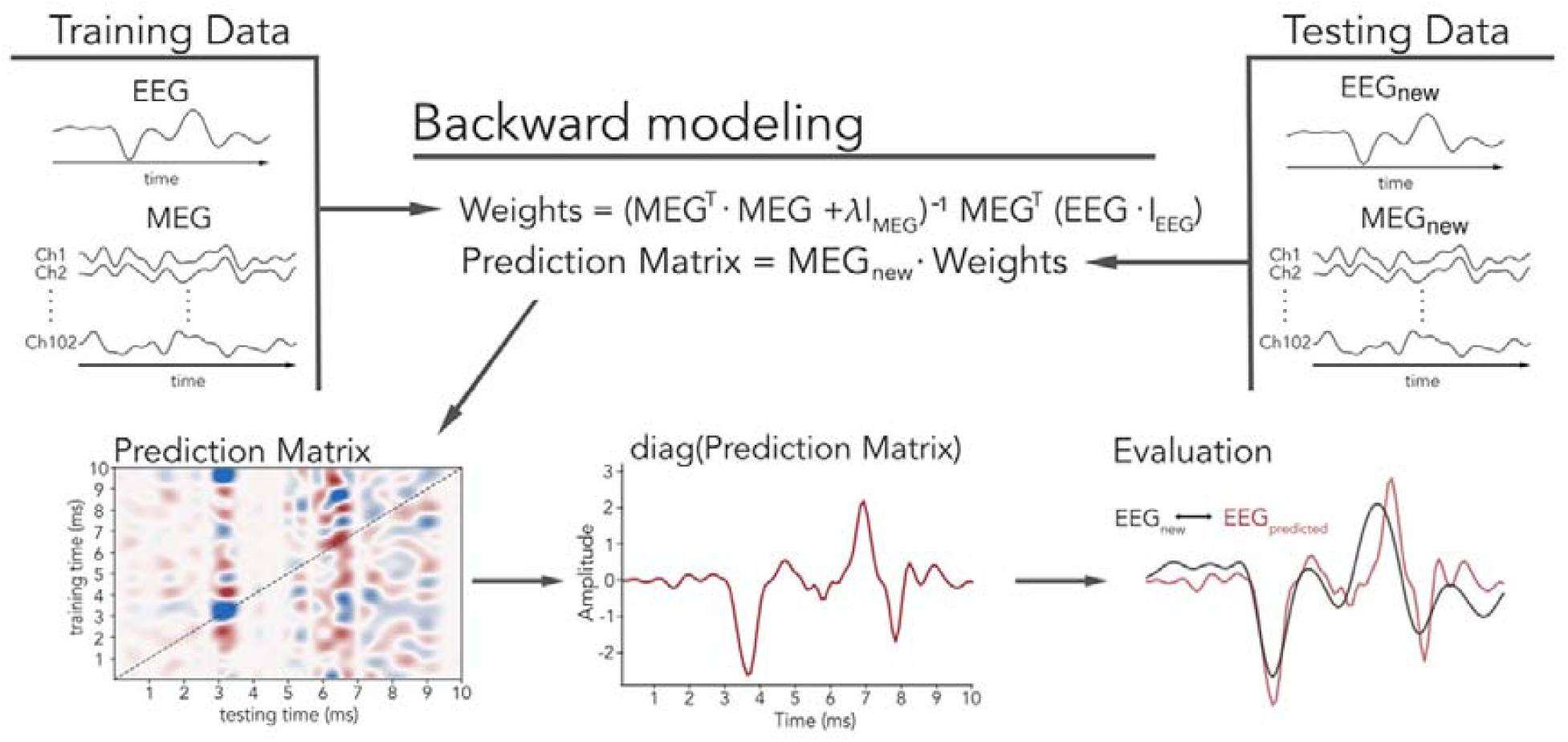
A backward modeling approach used to reconstruct auditory brainstem activity. MEG sensors were asserted as regressors for a concurrent electric ABR recording. The thereby generated weights were then used in a new measurement to reconstruct brainstem activity from the MEG recording. This activity was extracted by taking the diagonal from the prediction matrix. Here, the quality of the Prediction was evaluated by correlating the reconstructed ABR with the actual signal evoked by a different stimulation rate (9Hz).

To rule out that the weights generated during training will not reconstruct an ABR-like response independent of the measured input data, a reconstruction from surrogate data was performed and compared to the original reconstruction. The surrogate data was created by shuffling all timepoints in the averaged signal on each MEG sensor 1000 times. For each shuffle an overall correlation coefficient between the predicted and the original signal was computed. Finally, the correlation coefficients were z-transformed, averaged, re-transformed (Corey, Dunlap, & Burke, 1998) and compared to the correlation coefficient obtained from the real data using a Fisher r-to-z test.

#### 3.4.3 Source analysis

The structural MR-images of 7 participants were acquired by a Siemens MAGNETOM TIM Trio 3-Tesla MRI with 12-channel and 32-channel head coils. The individual brains were brought into a common space by co-registering the individual brains, based on the three anatomical landmarks (nasion, left and right preauricular points), with a standard brain from the Montreal Neurological Institute (MNI, Montreal, Canada). For the remaining participants without structural MR image, an anatomical template image was warped to the individual head shape. Afterwards a single-shell head model (Nolte, 2003) was computed for each participant. As a source model, a grid with 1 cm resolution and 2982 voxels based on an MNI template brain was morphed into the brain volume of each participant. This allows group-level averaging and statistical analysis as all the grid points in the warped grid belong to the same brain region across subjects. As backward model coefficients are spatially not interpretable (Haufe et al. 2014), linear forward model coefficients were computed (on the data obtained from the 30 Hz stimulation rate) using the same specifications as in the computation of our backward model. The resulting forward weights were then applied to an independent measurement using a 9 Hz stimulation rate, to predict the activity on the MEG sensors. Common linearly constrained minimum variance (LCMV) beamformer spatial filters (Van Veen, Drongelen, Yuchtman, & Suzuki, 1997) were then computed on the preprocessed MEG data and applied separately on the predicted and measured MEG activity to project them into source space. In order to show that the modeled brainstem activity is also spatially related to subcortical regions we correlated the predicted and measured activity for each voxel over the first 10 ms after stimulus onset. The resulting correlation coefficients were then compared to the average correlation coefficients of a moving correlation window between the predicted brainstem activity and measured activity during a pre stimulus interval (−20 ms to −10 ms). A dependent samples t-test was used to compare the z-transformed average correlation from the prestimulus condition with the z-transformed real condition. A non-parametric Monte-Carlo randomization test was applied to control for multiple comparisons (Maris and Oostenveld 2007). The test statistic was repeated 10 000 times on data shuffled across conditions. The observed clusters were compared against the distribution obtained from the randomization procedure and were considered significant when their probability was below 5%. For visualization, source localizations were mapped onto glass brain surfaces using nilearn (Abraham et al. 2014).

#### 3.4.4 Reconstructing brainstem activity using subsets of trials

In order to show that the generated model weights can be used to reconstruct brainstem activity in everyday listening situations (e.g. when listening to continuous speech or music) it is necessary to show that the reconstruction works when only a fraction of the usually needed amount of data is available. This was operationalized by applying the model weights to the evoked potential generated by random subsets of trials (5000, 1000, 500, 100, 50) drawn from the MEG data of the testing set (9Hz) for each subject. Each prediction was then z-scored and compared with the measured evoked potential (10 000 trials) by calculating the mean squared error between both signals. In a similar procedure random subset of trials (5000, 1000, 500, 100, 50) were drawn from the EEG data set (9Hz), averaged, z-scored and compared to the true evoked brainstem potential (10 000 trials). These results were then further analyzed using a two-way repeated measures ANOVA with the factors predicted ABR (EEG/backward model) and numbers of used trials (5000, 1000, 500, 100, 50).

## 4. Results

### Reconstructed brainstem activity is highly correlated with measured electric auditory brainstem activity

In order to reconstruct auditory brainstem activity from the MEG we performed a ridge regression between the signal captured by the MEG and EEG to generate a spatial filter for each training time point **(Figure 1, Backward modeling)** that putatively is “tuned” to brainstem activity. Our long-term goal following this proof-of-concept is to expand this approach to model auditory brainstem activity also in more interesting and complex listening situations.

In this study these channel weights were applied to an independent ABR measurement (i.e. testing data set) with a different stimulus frequency to predict the electrical ABR via backward encoding from the whole-head MEG data. This modeled activity was then correlated with the actually measured electric ABR. The results show a strong correlation between the measured and reconstructed activity on a single-subject level (**Figure 3a**), with a median correlation coefficient of *r*= 0.7, *p* < 0.0001. In order to exclude the possibility that the generated weights will reproduce an ABR-like response independent of the measured signal, we randomized the MEG data by shuffling all timepoints in the averaged signal on each MEG sensor (1000 times) before applying the channel weights to reconstruct the ABR. This reconstruction from randomized data was then correlated to the measured EEG activity. Finally, the correlation coefficients resulting from this permutation approach were compared to the previously empirically obtained correlation coefficients by means of a Fisher r-to-z test. The results show, the reconstruction from the real data was more strongly correlated to the actually measured EEG signal in all participants (**Figure 3c**). The difference between the correlations derived from real and randomized data was significant in 13 out of 14 participants. As the backward modelling did not work for subject 14 it was removed from all subsequent analysis.

**Figure 2:**
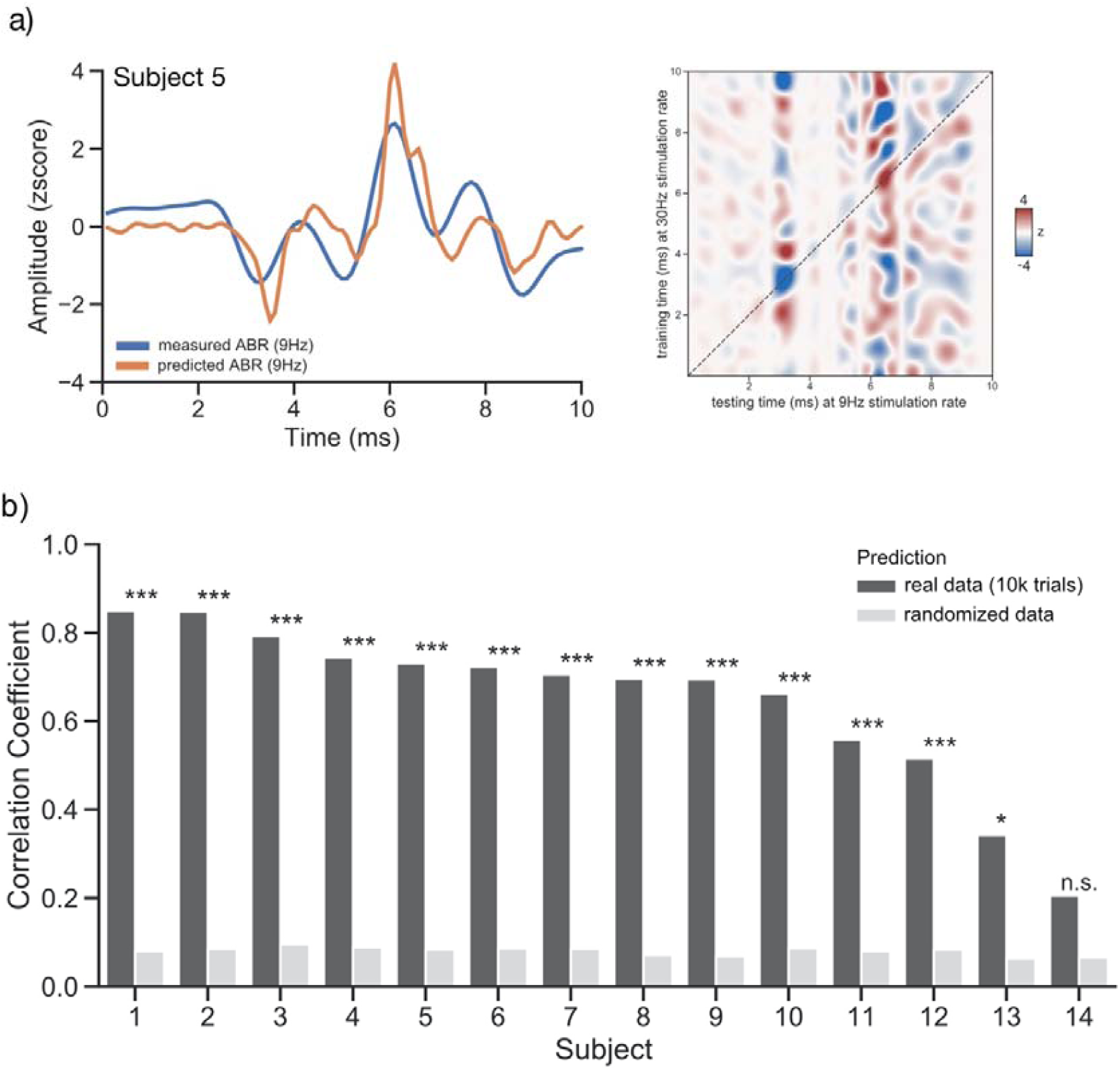
(a) The reconstructed brainstem activity (orange) for one representative subject (5; other subjects depicted in Figure S1) based on precomputed backward models trained on an independent dataset (30Hz stimulation rate), nicely tracks the actually measured brainstem activity in another independent measurement (9Hz; blue). The predicted ABR was extracted from the Temporal generalization matrix, marked by the dashed diagonal line. (b) Correlation coefficients on single subject level between the individually measured signal and its prediction (black) and between randomized signal and its prediction (grey). Significant differences between the prediction on the real and randomized data are indicated by asterisks.

**Figure 3:**
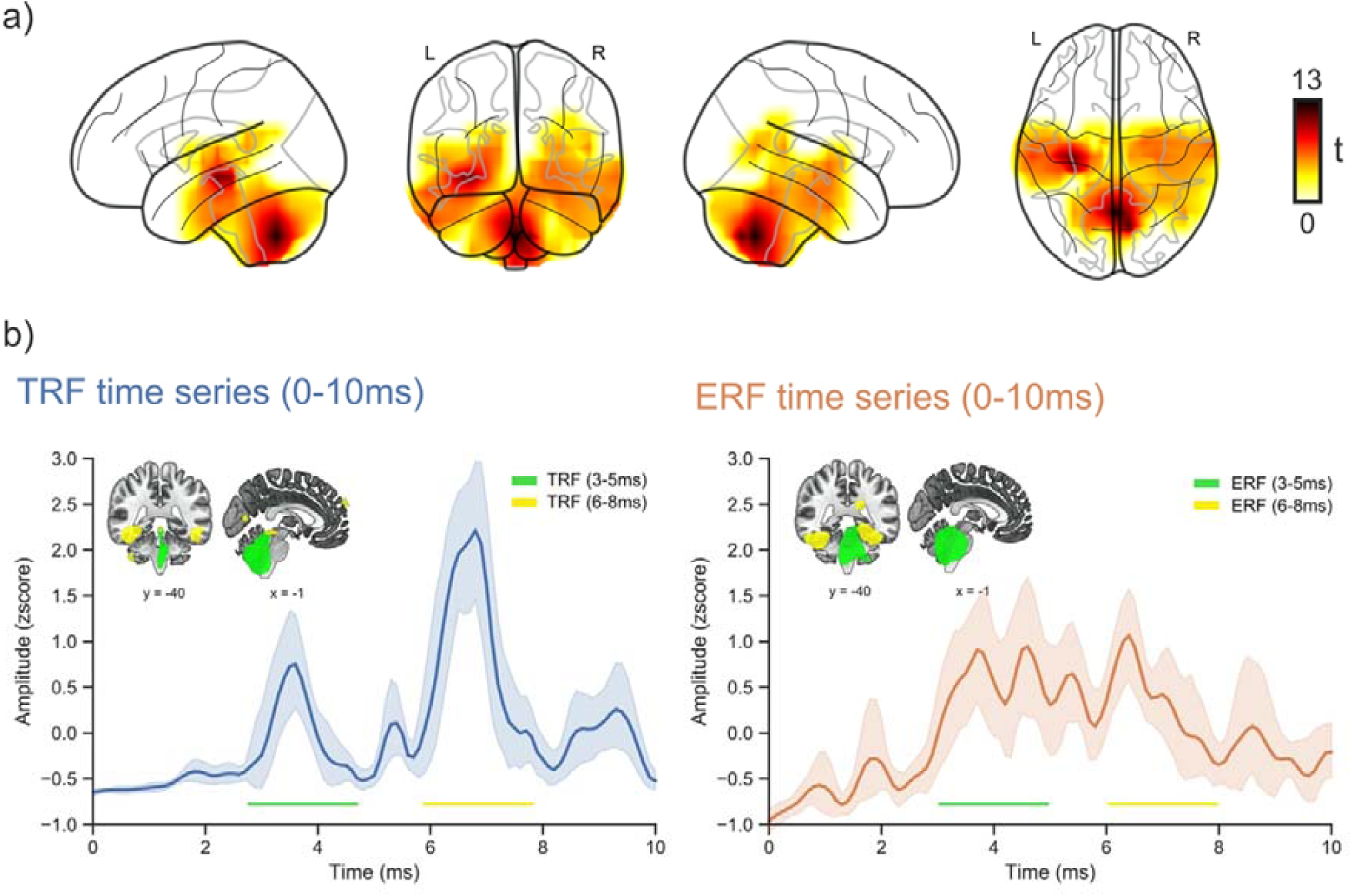
(a) Significant correlation between measured and predicted activity for 9Hz stimulation rate. The cluster stretches over subcortical and medial temporal regions crossing the auditory pathway. (b) The average TRF (blue) across all subjects and virtual channels (RMS) revealed two distinct peaks (3-5 ms/6-8 ms). The individually reconstructed sources (LCMV) across subjects for the modelled (blue) and measured response (orange) for these time courses showed that activity peaked between 3-5 ms (green) at locations in and close to the brainstem (observed and predicted). Activity between 6-8 ms (yellow) peaked at higher brainstem (predicted) and medial-temporal areas (observed and predicted). Activity was masked with an 80% threshold.

Apart from the high correlation between the reconstructed ABR (the diagonal of the temporal generalization matrix) and the measured electric ABR **(Figure 1, Evaluation)**, distinct off-diagonal patterns can be noted on the temporal generalization matrix in **Figure 2b**. This time generalized “activation” for each testing/training time point **(Figure 1, Signal extraction)** appears to be especially strong around the time course of wave V. This points towards a broader activation of structures along the ascending auditory pathway contributing to wave V than conventionally assumed (e.g. lateral lemniscus/inferior colliculi). While rarely explicitly reported a broad activation is actually in line with observations from invasive recordings indicating that many structures in the auditory brainstem are simultaneously activated by an acoustical stimulus (for a review; Biacabe et al. 2001).

### Model weights capture activity generated by subcortical regions

In order to estimate whether the model weights mainly capture activity generated by subcortical regions, forward model coefficients were computed using the same specifications as in the computation of the backward model. This was implemented to ensure a spatial interpretability of the coefficients in a neurophysiological manner. The forward weights were then used to predict subcortical activity at MEG sensors following a (9Hz) stimulation rate. Beamformer (LCMV) spatial filters were then computed on the preprocessed MEG data and applied separately on the predicted and measured MEG activity to project them into source space. The evoked activity at the virtual sensors for the predicted and measured activity were afterwards correlated across the time course of the ABR (0-10 ms). The resulting correlation coefficients were then compared to the average correlation coefficients of a moving correlation window between the predicted brainstem activity and measured activity during a pre-stimulus interval (−20 ms to −10 ms). A cluster-based permutation test revealed a difference between the correlation coefficients of the “real” and “pre-stimulus” condition (p<0.0001) in one positive cluster **(Figure 3a)**. The cluster stretches along subcortical and medial temporal regions crossing the auditory pathway with the highest correlations around the brainstem and cerebellum.

### Model weights show a progression of source localized activity along the auditory pathway

We demonstrated that measured and modelled auditory brainstem activity was highly correlated in subcortical and medial temporal areas crossing the auditory pathway. But does the modelled brainstem activity also reflect the expected progression of auditory evoked activity during the first 10 ms along the auditory hierarchy from lower to higher order processing areas? In order to investigate whether we could observe such a transition of auditory evoked activity we first averaged (RMS) the TRF’s/ERF’s across all virtual channels per subject. The resulting individual global TRF’s/ERF’s were then averaged across subjects (**Figure 3b**). This revealed two main distinct peaks at 3-5 ms and 6-8 ms in the TRF time series **(blue)**. When reconstructing the sources for the two time courses separately for the measured and predicted activity using LCMV beamformers we noticed that the predicted and measured activity between 3 and 5 ms (**Figure 3b, green**) is distributed focally at regions in and close to the brainstem (thresholded at 80%). In contrast, the modelled activity for the later time interval between 6 and 8 ms (**Figure 3b, yellow**) was distributed more widely with activity at upper brainstem (predicted), but also medial temporal areas between brainstem and primary auditory cortex (predicted and measured; thresholded at 80%). Descriptively, this shows a spread of neural activity from lower to higher order processing areas both in the actually measured and predicted signals.

### Brainstem activity reconstructable in situations with a low signal-to-noise ratio

So far it was shown that auditory brainstem activity can be reconstructed using backward encoding models and that the associated forward model weights mainly capture activity generated by subcortical regions. However, to show that the proposed modelling approach can also be used in situations with a lower signal-to-noise ratio it is necessary to show that the model weights can be used to reconstruct brainstem activity when only a fraction of the usually needed amount of data is available.

In order to show this the model weights were applied to the evoked potential generated by random subsets of trials (EEG data (9Hz), 5000, 1000, 500, 100, 50) drawn from the MEG data of the testing set (9Hz) for each subject. Afterwards the mean squared error between each subset and the true evoked response (EEG data (9Hz), 10 000 trials) was computed. This procedure was repeated 1000 times. The average mean squared error for each subject is depicted in **Figure 4 (orange)**. Additionally, the mean squared error between the true evoked response (EEG data (9Hz), 10 000 trials) and the evoked potential generated by random subsets of trials (EEG data (9Hz), 5000, 1000, 500, 100, 50) drawn from the EEG data was computed **Figure 4 (green)**.

**Figure 4:**
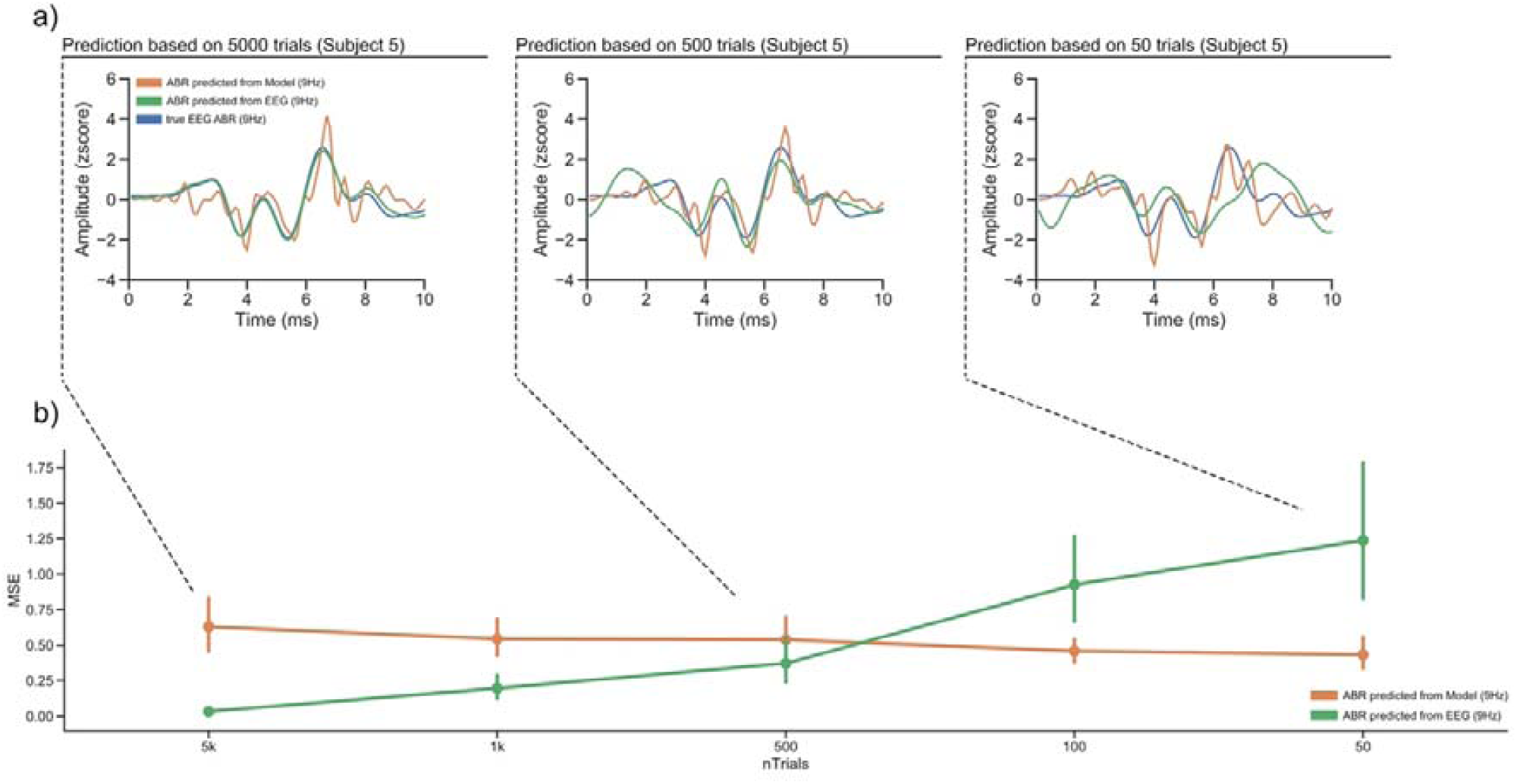
(a) Randomly selected subsets of trials in the testing dataset (9Hz) were used to reconstruct an ABR here shown for one representative subject (5; other subjects depicted in Figure S2). Brainstem activity predicted based on a subset of trials using the backward model and the associated signal measured using an EEG electrode were compared to the true evoked response elicited by 10 000 trials and measured via EEG. (b) Mean squared errors between predicted ABR (EEG (green)/the backward model (orange)) and the true response over all subjects show that the reconstruction of auditory brainstem activity is still possible in situations with a lower signal-to-noise ratio.

The results of a two-way repeated measures ANOVA revealed that there was a significant main effect for the amount of presented trials on the mean squared error (F(4, 48)=21.693, *p*=0.00006, η_*p*_^*2*^=0.644). In situations when more trials were presented the actual ABR was better predicted as in situations where fewer trials were presented, shown by a lower mean squared error. There was no significant main effect on the factor used to predict an ABR (eeg/backward model) on the mean squared error (F(1, 11)=0.301, *p*=0.593, η_*p*_^*2*^=0.024). However, there was a significant interaction effect between the factor used to predict an ABR (eeg/backward model) and the presented amount of trials (F(4, 48)=27.588, *p*=0.00001, η_*p*_^*2*^=0.697) such that the difference between the ABR evoked with subsets of the EEG data compared to the ABR evoked by the full dataset increases when fewer trials are used to compute the evoked potential. This was not the case for the modeled ABR. Here the mean squared error between the average individual reconstructions and the associated measured EEG responses for each subset was stable even in situations where only 50 trials were used to compute an ABR. This shows that the model weights can be used to reconstruct an ABR in situations with a low signal-to-noise ratio.

## 5. Discussion

In this proof of concept study, we introduce a new approach to capture auditory brainstem activity using a combination of EEG and MEG. Since the presented approach incorporates a (time-) generalization step by applying it to a new dataset, it holds promise to enable the modeling of brainstem activity also in more naturalistic auditory scenarios (e.g. listening to continuous speech or music). So far this has been problematic, as deriving auditory brainstem activity requires averaging over thousands of simple sounds to uncover the signal produced by brainstem regions, resulting in a highly artificial measurement environment. Our approach circumvents this issue by using the signal obtained during a MEG recording as regressors for a concurrent electric ABR measurement. Through this backward encoding approach channel weights that are responsive to activity from putative auditory brainstem structures are generated **(Figure 1)**. We propose that these channel weights can be applied to any other data using the same sensor coverage. Analogous to classification based decoding approaches (King & Dehaene, 2014), we assume that the temporal generalization matrix resulting from the application of the channel weights to a new dataset can be interpreted as the presence of an activation pattern generated during a training period (e.g. the pattern contributing to a specific ABR wave) at a corresponding testing time (e.g. cue target period in selective attention tasks).

Here, we validated this approach by using these weights to reconstruct the expected brainstem activity from the magnetometers in another ABR measurement with a different stimulation rate (9 Hz). This reconstructed response was then compared to the measured response showing that both were highly correlated. This validation marks a first step prior to using this method to model brainstem activity within complex and natural auditory scenarios (e.g. continuous speech, music). On top of that we showed that the modeled brainstem activity is also spatially related to subcortical regions. This was accomplished by calculating new model weights in a forward direction and correlating the source reconstructed prediction with the actually measured MEG activity. The resulting effect was strongest in subcortical regions in and close to the brainstem, corroborating our assumption that our approach provides a means of capturing activity from subcortical areas along the auditory hierarchy. This assumption was further supported by showing that measured and modelled activity transitioned from deeper areas within an early time interval (3-5 ms) to higher order areas during a later time interval (6-8 ms). This is in line with invasive recordings showing that the main generators and pathways associated with earlier deflections (e.g. wave III) are situated deeper than those of later deflections (e.g. wave V) (Wada, 2012). Additionally, we observed activity in medial temporal regions between brainstem and primary auditory cortical areas from 6 to 8 ms in the predicted and measured signals. This is in agreement with results from human invasive recordings that show an initial afferent thalamocortical volley with an onset ∼9 ms at posteromedial parts of Heschl’s Gyrus (Liégeois-Chauvel, Musolino & Chauvel, 1991; Brugge et al., 2008). While these observations are intriguing and in agreement with previous invasive recordings, they should still be interpreted with caution, as both measured and predicted activity stretch over areas beyond the nuclei commonly associated with activity on the auditory pathway. In part, this lack of accuracy may be attributed to the low spatial acuity of MEG compared to invasive or MRI recordings.

However, as the backward models are trained to predict a widely accepted surface-recordable brainstem response it offers the possibility to omit the source reconstruction procedure entirely in the future. This could be of interest in situations where only EEG is available and source localization procedures are overall more challenging compared to MEG.

Additionally, we showed that the proposed modeling procedure can be applied to situations where the signal-to-noise ratio is low (e.g. when only a few trials are available). This marks an important first step prior to using this method to model brainstem activity within more complex and natural auditory scenarios (e.g. continuous speech, music). Here, a combination with alternative approaches may be interesting. For example, Maddox and Lee (2018) succeeded in measuring brainstem activity following continuous speech by asserting the rectified speech audio signal as regressor for their EEG recording. They found a distinct peak at ∼6 ms that was highly correlated with their click evoked ABR. Evoked activity during that time course is commonly associated with Wave V of the ABR (putatively lateral lemniscus/inferior colliculi (Møller & Jannetta, 1985)). Similarly to Maddox and Lee (2018) we noted a peak at the time course of Wave V, but also found in most subjects (even in situations with an unfavorable signal-to-noise ratio) a peak at the time course of Wave III (putatively mainly cochlear nucleus/ superior olivary complex (Møller & Jannetta, 1985; Møller, Jho, Yokota, & Jannetta, 1995). This is an important addition, as using the here proposed approach as a “preprocessing” procedure to tune a measurement device (EEG/MEG) to different subcortical areas prior to deriving TRF’s for a rectified audio signal (e.g. continuous speech, music) would potentially allow researchers to gather more insight on the involvement of deeper subcortical regions during auditory processing.

Using a similar approach as Maddox and Lee (2018), by cross-correlating the fundamental waveform of continuous speech with the recorded EEG signal, Forte et al. (2017) found a distinct peak at ∼9 ms. This evoked activity was modulated by attention, leading the authors to suggest a mechanism of selective attention at the level of the brainstem. However, with occurrences of early auditory cortical activity measured directly at posteromedial parts of Heschl’s Gyrus ∼9-10 ms at least some thalamocortical contributions are likely (Liégeois-Chauvel, Musolino & Chauvel, 1991; Brugge et al., 2008). In any case also this approach could be combined with ours, which could improve the rather small effect sizes for the reported attention effect (*r* < 0.06).

However, the TRF (Maddox & Lee, 2018) as well as the cross-correlation approach (Forte, Etard & Reichenbach, 2017) require a continuous sound stimulation to measure auditory brainstem activity. In principle our trained backward models are not dependent on a continuous sound stimulation and could also be applied to silent periods (e.g. a cue target period, sound omission periods; e.g. Frey et al. 2014; Demarchi, Sanchez & Weisz, 2019). One of the potentially exciting perspectives would be to estimate such ongoing brainstem activity during cognitive tasks. However, future studies ideally complemented by invasive works are necessary to assess how versatile our proposed approach is.

## 6. Conclusion

In this study, we used a combination of MEG and EEG to create a backward encoding model that can be used to reconstruct auditory brainstem activity. We validated this approach by using the weights generated during an ABR measurement with a 30 Hz stimulation rate to predict the expected EEG ABR activity from the MEG in another measurement with a different stimulation rate (9 Hz). By showing that the expected and modeled response are alike (even in situations with an unfavorable signal-to-noise ratio) we established the important proof-of-concept that backward encoding models can be generalized to an independent data set to predict an ABR with high accuracy. The power of (time-)generalizing the backward encoding model to different data sets (using the same sensor coverage), could improve the outcomes and interpretation of existing efforts (Maddox & Lee, 2018; Forte et al. 2017) but could also in principle enable the modeling of auditory brainstem activity in other more interesting listening situations. The further development of this approach may enhance our understanding of auditory processing by making the contributions of subcortical structures measurable in a non-invasive fashion.

## 7. Acknowledgements

We thank Maja Serman and Ronny Hanneman for discussion. This research was supported by the Sivantos GmbH.

## 10. Supplementary Materials

**Figure S1:**
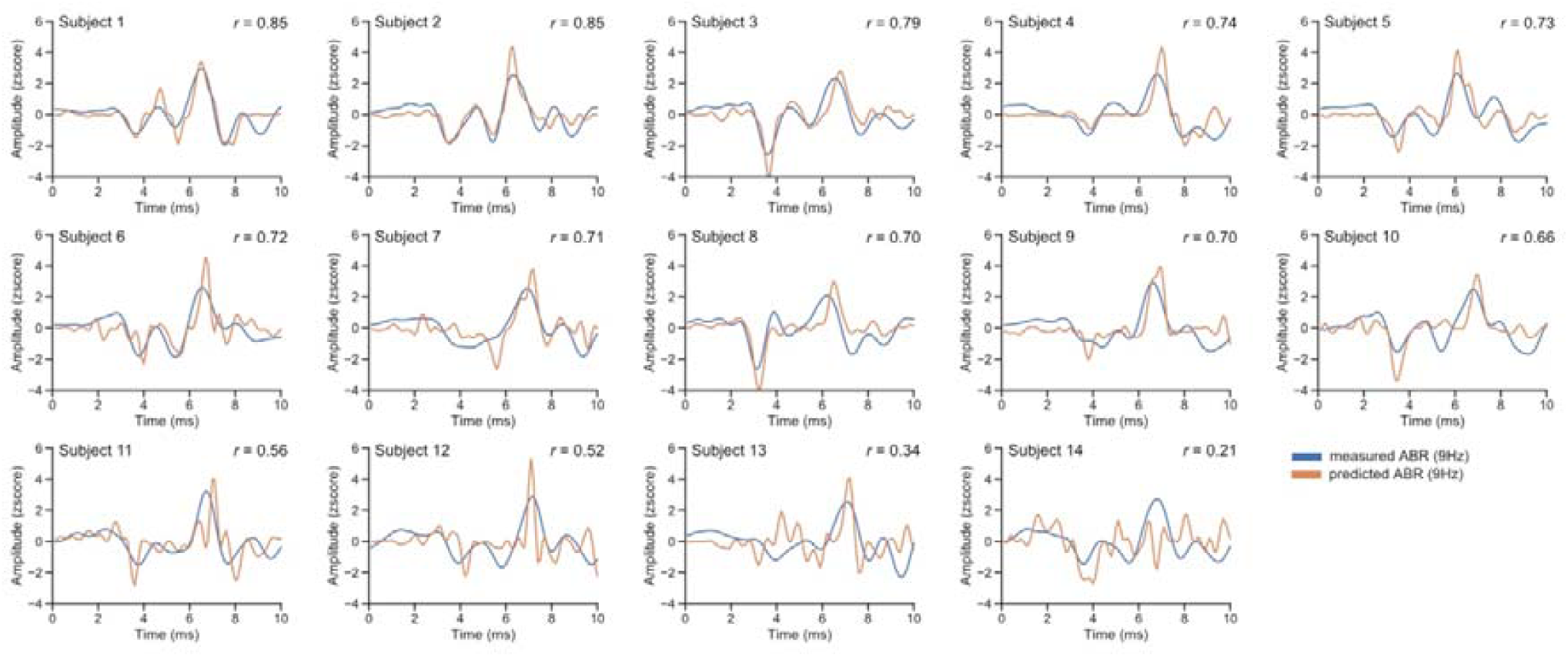
The reconstructed brainstem activity (orange) for each subject based on precomputed backward models trained on an independent dataset (30Hz stimulation rate), tracks the actually measured brainstem activity in another independent measurement (9Hz; blue). The predicted and measured activity was correlated significantly in 13 out of 14 participants.

**Figure S2:**
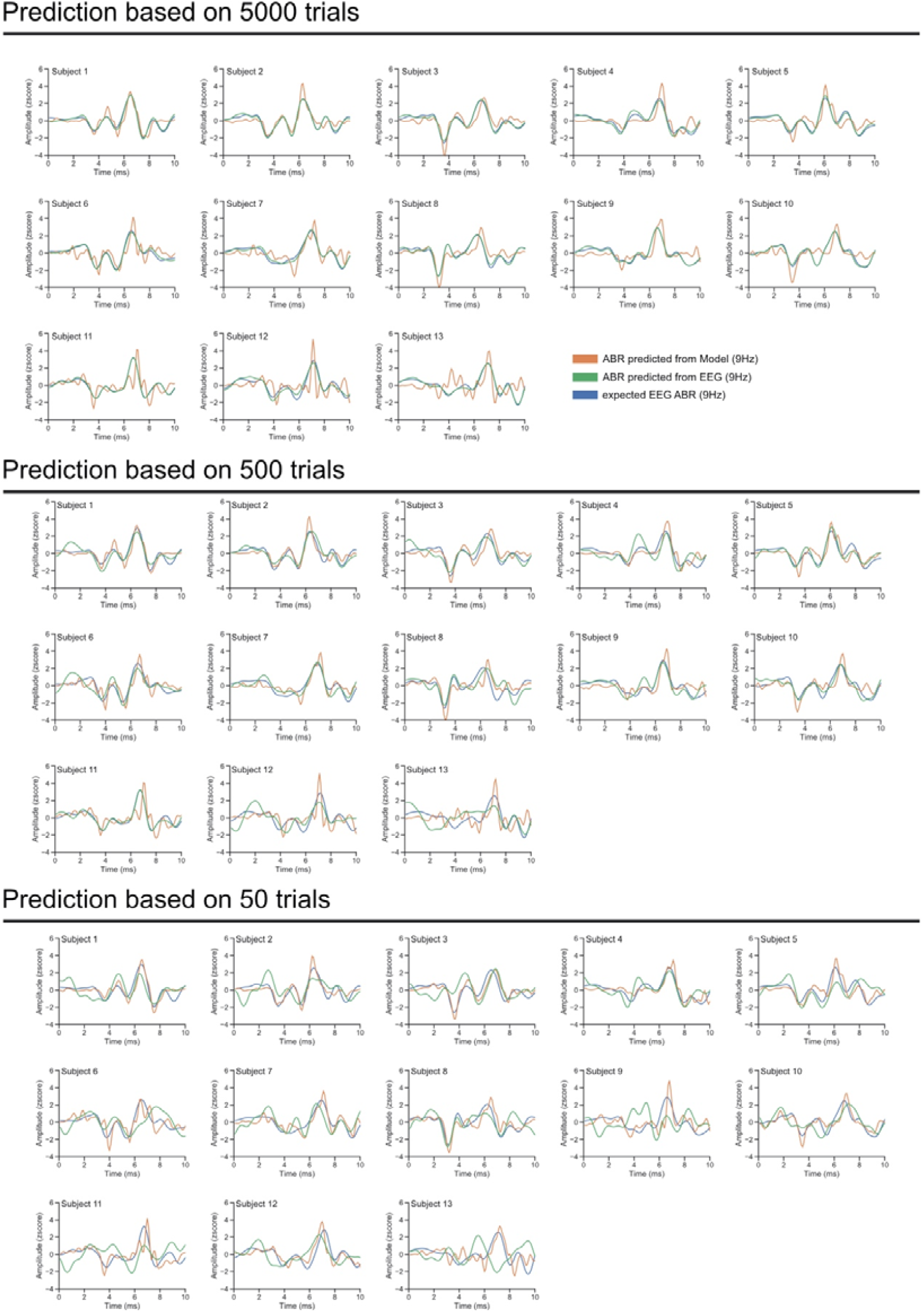
The reconstructed brainstem activity (orange) and measured brainstem activity (green) for only a subset of trials compared to the true brainstem activity (blue) for all presented trials.

